# Predicting complex phenotypes using multi-omics data in maize

**DOI:** 10.1101/2025.09.30.679283

**Authors:** M Creach, B Webster, L Newton, J Turkus, JC Schnable, A Thompson, R VanBuren

## Abstract

Understanding and predicting complex traits in plants remains a fundamental challenge due to the emergent nature of most phenotypes and their dependence on genetic, regulatory, and environmental interactions. Accurate prediction of traits and identification of underlying genetic elements has broad applications for plant breeding, systems biology, and biotechnology. Here, we tested if multi-omic datasets could improve predictive accuracy of 129 diverse maize phenotypes across nine environments using genomic markers, field based transcriptomic data from two locations, and drone-derived phenomic data of vegetative indices. We trained and compared linear (rrBLUP) and nonlinear (support vector regression) models using single- and multi-omics inputs. Multi-omics models consistently outperformed single-omics models for most traits, with genomic and transcriptomic inputs contributing distinct biological features. Phenomic features alone yielded the lowest predictive power but improved predictions for specific trait categories like root architecture. Transcriptomic datasets enabled cross-environment prediction, demonstrating that gene expression patterns from one field site could accurately predict traits measured in another. Environment-specific expression of benchmark flowering time genes highlighted the value of transcriptomics in capturing genotype-by-environment (G×E) interactions not detectable through genomic data alone. These findings demonstrate that integrating transcriptomic and phenomic data with genotypes enhances trait prediction, improves model generalizability across environments, and provides deeper insight into the genetic and regulatory architecture of agriculturally important traits in maize.

## Introduction

Predicting complex traits remains a fundamental challenge in biology because phenotypes emerge from the interactions between genetic variation, dynamic biological processes, and environmental conditions. Phenotypes are shaped not only by environmental factors but also by emergent properties of gene networks and cellular processes, where many genes interact in complex and context-dependent ways. Understanding how genetic variation translates into phenotypic variation has broad applications in many fields including plant breeding, systems and synthetic biology, evolutionary biology and biotechnology.

Numerous statistical and machine learning approaches have been developed to predict phenotypes from genotypic and other increasingly high-dimensional biological datasets. Genomic prediction is now a cornerstone in biology, where a range of modeling frameworks are used to link genome-wide data to trait outcomes. Linear models such as rrBLUP remain popular due to their stability and strong performance for highly polygenic traits ^1,2^. However, no single approach is optimal for every context. Bayesian models like BayesA, BayesB, and Bayesian LASSO allow marker-specific effect sizes and often outperform linear models for traits with large-effect loci ^3,4^. Nonlinear models such as random forest, support vector regression, and deep learning can capture complex patterns in high-dimensional data, including interactions among features and non-additive effects ^5,6^. Model performance depends on factors such as population structure, heritability, environmental variability, and training set size in both plants and animals ^7^.

Efforts to improve predictive accuracy have increasingly focused on integrating additional data types that reflect biological processes in real time. For example, transcriptomic and phenomic datasets have improved predictions for flowering time, abiotic stress tolerance, and developmental traits by capturing dynamic responses that are not encoded in static genomic markers ^8–10^. This improvement is not strictly context dependent, as gene expression data collected from maize seedlings improved prediction of adult phenotypes and identified more known flowering-time genes than models based solely on genotypes ^11^. Other studies have shown similar improvements for disease resistance in wheat ^12^ and flowering time in Arabidopsis ^13^. However, these studies typically evaluate a small number of traits, often with relatively simple genetic architectures and rely on limited omics sampling across environments, leaving open questions about broader generalizability and environmental robustness.

In this study, we tested how combining diverse omics data types with both linear and nonlinear models impacts predictive accuracy for 129 agronomic traits in maize. We used gene expression data collected from field-grown plants at two locations, under the hypothesis that transcriptomic and phenomic data reflect condition-specific biological responses that can capture trait-relevant variation beyond genomic markers alone. By training and testing models using genomic, transcriptomic, and image-based phenomic inputs, we evaluated the relative contribution of each data type. Our results show that combining these data types improves prediction accuracy for most traits and that the additional omics inputs provide complementary, nonredundant information. These findings suggest that integrating biologically rich datasets across environments can help explore the basis of complex trait variation, enhance predictive modeling for crop improvement, and uncover additional loci for targeted breeding or functional analysis.

## Results

### Building a multi-omics framework for predictive trait modeling

We set out to test whether integrating high-dimensional omics datasets could predict complex traits as well as or better than using genomic data alone. This was based on our hypothesis that transcriptomic and phenomic data reflect real-time biological and environmental responses that may capture additional predictive information. We chose maize as a model to test this because of its immense wealth of phenotypic and agronomic data collected across diverse environments, exceptional genetic diversity, and the availability of large-scale, systems-level datasets including genomics, transcriptomics, and high-throughput phenotyping. We assembled three types of input features from ∼750 accessions in the maize Wisconsin Diversity Panel and evaluated their predictive power both separately and in combination. These input datasets included: genomic data (G), consisting of SNP-based markers; transcriptomic data (T), consisting of gene expression profiles; and phenomic data (P), consisting of image-derived vegetative indices from drone surveys. We built predictive models for 129 diverse maize traits ^14^ using each data type alone (single-omics models) and using combined datasets (multi-omics models). We compared a standard linear modeling approach (ridge-regression G-BLUP, implemented as rrBLUP) against a nonlinear machine learning approach (Support Vector Regression, SVR) to see if nonlinear patterns in the data would improve accuracy. Model training and testing followed a five-fold cross-validation scheme. The ∼750 panel genotypes were randomly divided into five folds (each ∼20% of genotypes), and in each iteration one fold was held out for testing while the model was trained on the other four folds. This ensured each genotype and all its trait values was predicted exactly once with no data leakage. We repeated this process for every combination of input data (G, T, P, and selected multi-omics combinations such as G+T and G+T+P) and for both algorithms, yielding a total of 1,290 trained models (Figure 1). Prediction performance was evaluated by the Pearson Correlation Coefficient (PCC) between predicted and observed trait values in the test sets.

**Figure 1.**
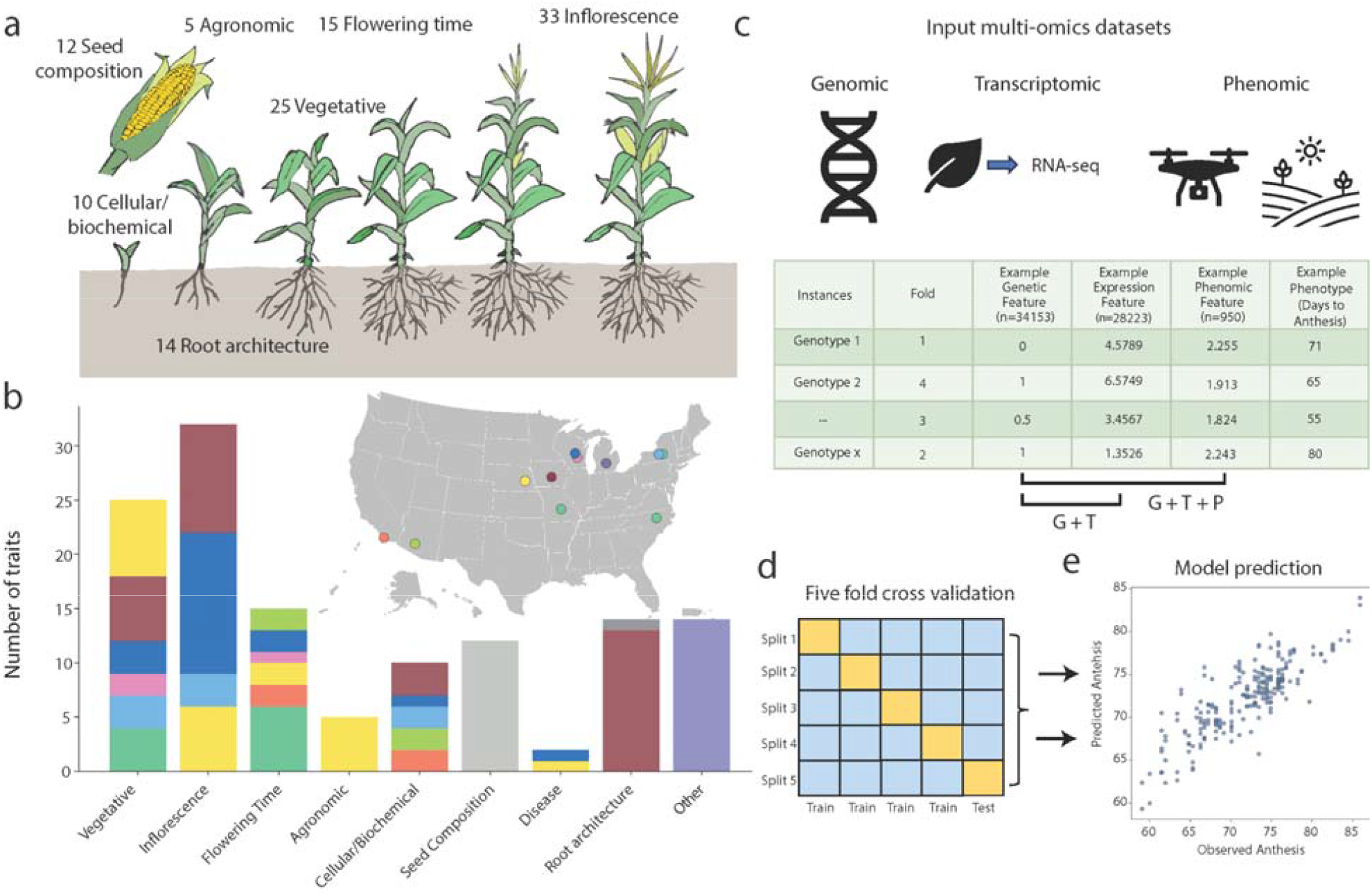
Overview of input datasets and modeling framework for multi-omic trait prediction in maize. (a) Traits included in the predictive modeling framework span diverse categories including vegetative morphology, inflorescence structure, flowering time, root architecture, seed composition, agronomic traits, disease resistance, and cellular/biochemical traits. (b) Location and distribution of the 129 phenotypes used for predictive modeling. Data collection sites are shown on the map, with bar plot colors corresponding to the number of phenotypes in each category collected at the respective location. (c) Overview of input features derived from genomic (n = 34,153 SNPs), transcriptomic (n = 28,223 gene expression values), and phenomic (n = 950 vegetative indices) datasets. Models were trained using single-omics datasets or combinations. (d) Predictive models were trained using five-fold cross-validation, with each fold iteratively serving as a test set while the remainder were used for training. (e) Example model prediction of anthesis date using transcriptomic data. Observed days to anthesis (X-axis) are plotted against predicted values (Y-axis) for each accession in the maize Wisconsin Diversity panel.

The genomic dataset (G) was derived from whole genome resequencing data of the Wisconsin Diversity Panel ^15^ filtered to a set of 34,153 high-quality single nucleotide polymorphisms (SNPs) by removing heterozygous sites and pruning markers in linkage disequilibrium. For the transcriptomic dataset (T), RNAseq data was collected from leaf tissue of the Wisconsin Diversity Panel collected in the field within a two-hour window around the time of flowering in Michigan and Nebraska field trials. RNAseq reads were aligned to the maize B73 V5 reference genome, and transcripts were filtered to remove any genes with low expression or no variance, resulting in a set of 28,223 genes. Transcript abundances were log-transformed and batch-corrected by state using PyCombat to account for technical differences and other artifacts, and the resulting normalized TPMs were used as transcriptomic features in all models. The phenomic dataset (P) comprises 950 vegetative indices extracted from the drone imagery of 12 flyovers of the Wisconsin Diversity Panel, collected from the same Michigan field plots used for RNA-seq.

To assess whether the omics inputs captured distinct sources of biological variation, we conducted principal component analyses (PCA) on each dataset. Genetic PCA using an expanded SNP set showed weak clustering by subpopulation, reflecting the extensive diversity and recombination within the Wisconsin Diversity Panel (Figure 2a). PCA of the log transformed expression data with no batch correction separated the samples by field site (e.g., Michigan vs Nebraska; Supplemental Figure 1). Batch correction removed technical artifacts between sites, and the first two principal components separated the samples by major genetic subgroups of stiff stalk and non-stiff stalk (Figure 2b). Phenomic PCA did not reveal strong subpopulation clustering, consistent with the expectation that image-derived features primarily reflect environment-responsive traits rather than genetic structure (Figure 2c). To assess how similar these datasets are quantitatively, we calculated pairwise PCCs of all input datasets. We observed a moderate correlation between genetic and transcriptomic data (PCC = 0.59), consistent with some degree of genetic control of transcriptomic variation, as expression patterns often exhibit moderate heritability within structured populations ^16–18^. In contrast, phenomic data showed minimal correlation with either genomic (PCC = 0.052) or transcriptomic (PCC = 0.12) inputs, suggesting that image-derived phenotypes capture orthogonal information or residual variation (Fig. 2d). These results indicate that each of our input datasets contains unique information about biological processes that could be utilized in a predictive model.

**Figure 2.**
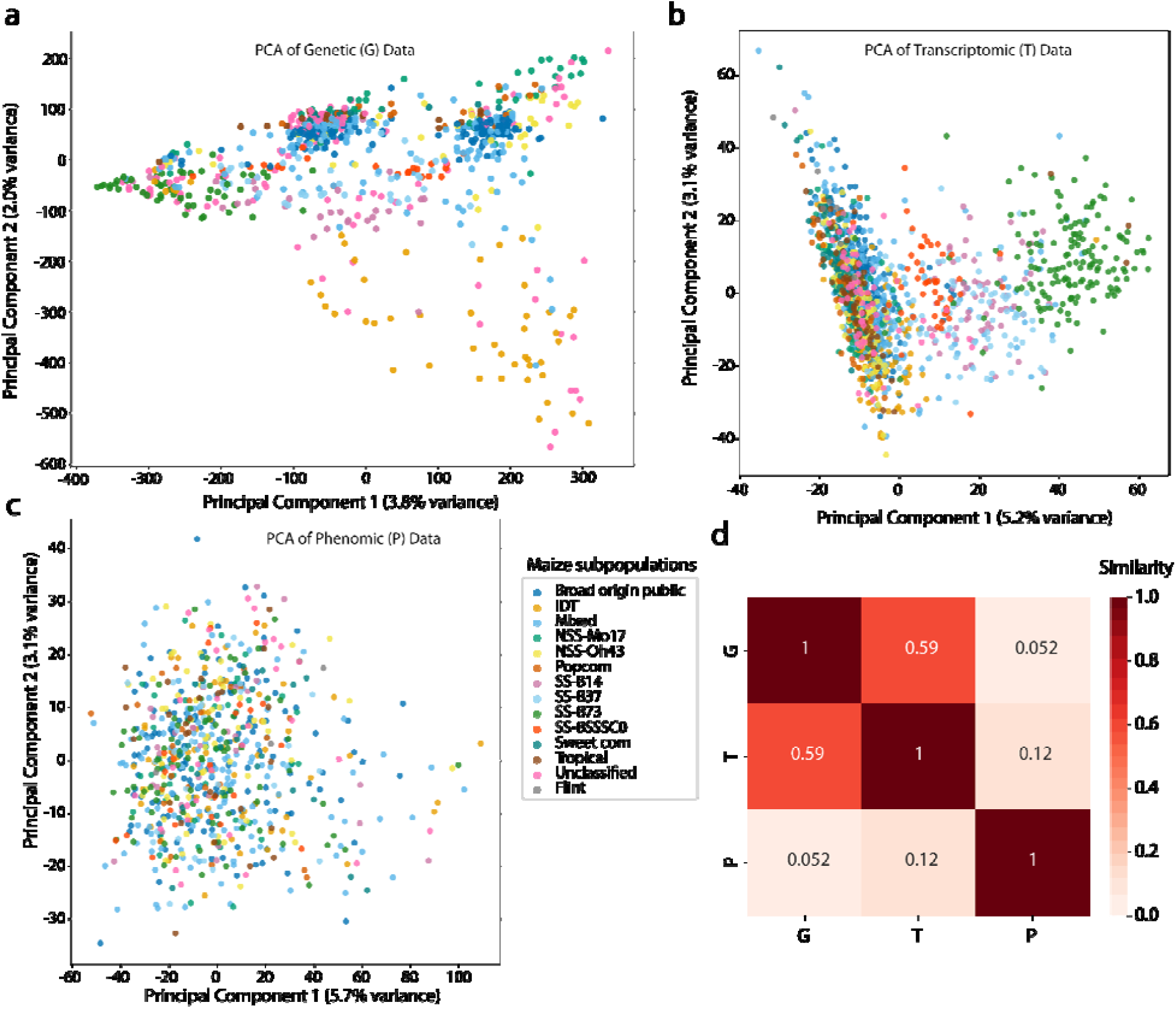
Dimensionality reduction and correlation of input datasets used for predictive modeling. Principal component analysis (PCA) of genomic (a), transcriptomic (b), and phenomic (c) datasets. The first two principal components are plotted for each dataset and colored by subpopulations within the Wisconsin Diversity Panel. Transcriptomic data was batch corrected and log2+1 transformed prior to analysis. (d) Pearson correlation coefficient similarity between pairwise combinations of all three input datasets.

### Model performance using different omics inputs over 129 complex phenotypes

To test the predictive power of different omics datasets, we trained predictive models for all 129 maize traits using rrBLUP and SVR with genomic, transcriptomic, phenomic, and combined multi-omics data. We tested rrBLUP, a linear mixed model that is commonly used in genomic prediction, because it assumes additive genetic effects and models the relationship between genetic markers and traits as linear. We tested SVR to evaluate whether a nonlinear machine learning algorithm would capture more complex interactions among features and have a higher predictive performance. The 129 phenotypes encompass a broad range of trait categories, including flowering time, agronomic performance, vegetative growth, seed composition, root architecture, biochemical and cellular features (Figure 1a, Supplemental Table 2). These traits were measured in the Wisconsin Diversity Panel across multiple field sites, locations, and growing seasons (Figure 1b). Thus, the omics data were used to predict phenotypes from other field sites and years beyond the original experiment in which they were collected. To establish a baseline for our predictions, we used the first 75 principal components of the genetic data as a proxy for population structure in all models. For each trait, model performance was measured using the Pearson correlation coefficient between the actual and predicted phenotype values (Figure 1e).

Multi-omics models have higher mean predictive accuracy than the single-omics models (Fig 3a). Models built using rrBLUP generally outperformed those using SVR, although the differences in performance distributions were not statistically significant as determined by ANOVA followed by Tukey’s HSD test. Models that combined genomic and transcriptomic data (G +T) often achieved higher accuracy than the baseline genomic models or models with just transcriptomic data. Incorporating phenomic data to the multi-omics models did not consistently improve the predictive performance and in some cases even lowered the performance for certain traits. Models using only phenomic data had the lowest predictive accuracy overall. This is expected as vegetative indices derived from drone imagery capture limited information about underlying physiological or developmental processes, and are especially uninformative for traits like seed composition that are determined later in the life cycle or in non-vegetative tissues.

**Figure 3.**
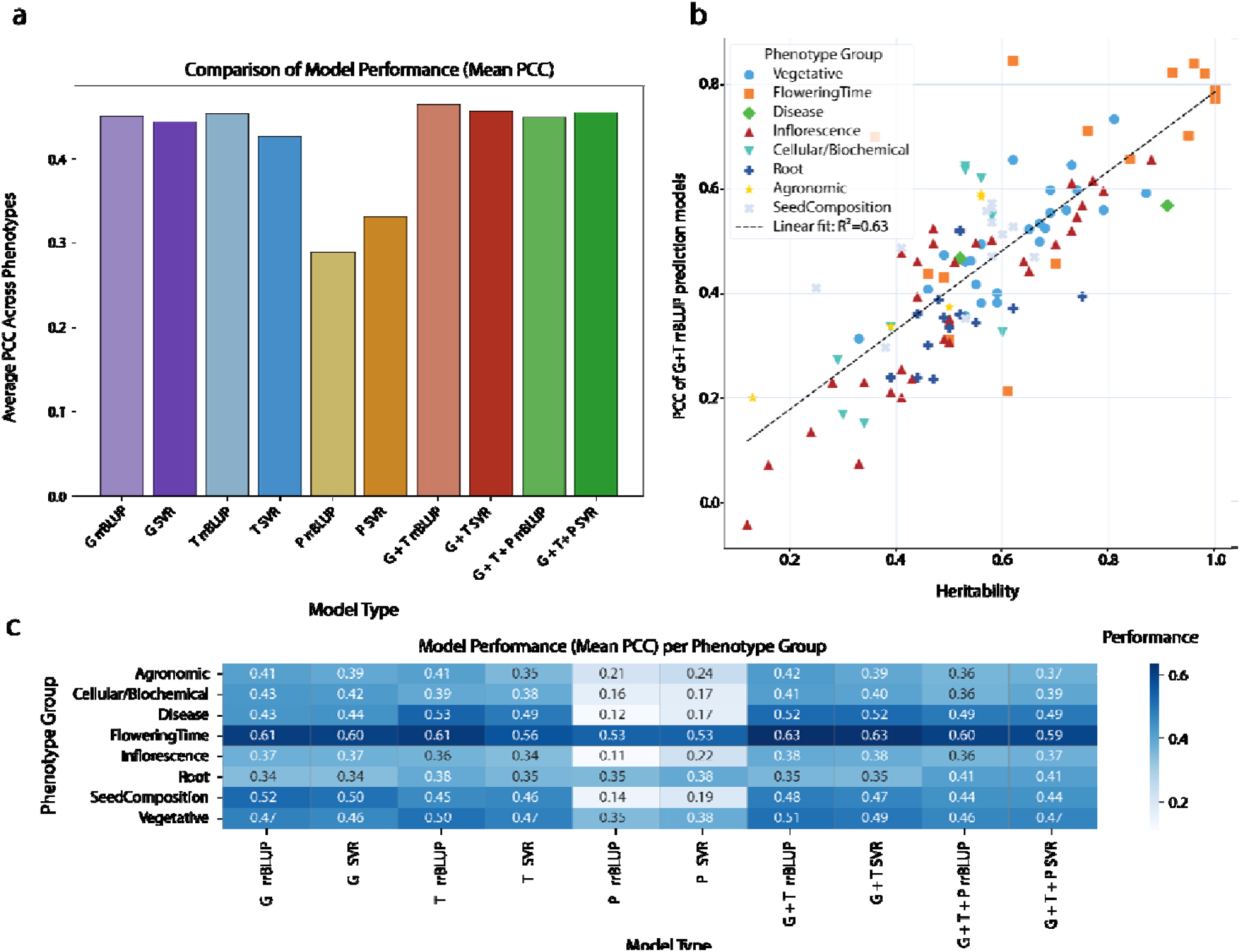
Predictive performance of different omics models across diverse maize phenotypes. (a) Average model performance (Pearson correlation coefficient, PCC) across all 129 traits for each omics input type and algorithm. (b) Relationship between trait heritability and model performance using the rrBLUP model with genomic and transcriptomic data (G+T). Each point represents a single trait, colored by phenotype category (R^2^=0.63, p=0.871e-25). (c) Mean predictive accuracy (PCC) per phenotype group across different model types.

We observed a positive relationship between the narrow-sense heritability of each trait and the predictive accuracy (Figure 3b), indicating that traits with higher heritability tend to be more accurately predicted. For example, flowering time traits, which are highly heritable ^19^, consistently achieved strong model performance. In contrast, root traits, which are generally more environmentally influenced and less heritable ^20^, showed lower predictive accuracy, regardless of the input data or algorithm. This trend underscores the importance of underlying genetic architecture in determining the success of predictive modeling.

To better understand how model inputs affect trait-specific predictions, we examined the mean PCC across the phenotype groups as defined by Mural (2022). No one model type or input dataset performed the best across all phenotypes, and optimal prediction is trait specific (Figure 3c). There was a general increase in model performance with the inclusion of transcriptomic data but this was not universal (Figure 3c). Flowering time related traits have the overall highest PCCs, and models perform best when using both genetic and transcriptomic data, regardless of the model type (mean PCC = 0.63 for both SVR and rrBLUP). Roots traits have the highest mean predictive accuracy when using multi-omics models that combine genomic, transcriptomic, and phenomic data (mean PCC = 0.41 for both SVR and rrBLUP). While no single input type performed well on its own, the inclusion of phenomic data improved predictions for root traits, suggesting that above-ground vegetative indices such as growth rate, canopy structure, and biomass carry predictive signals related to below-ground architecture. In contrast, seed composition traits are best predicted using rrBLUP models with genomic data (PCC= 0.52), likely due to their strong genetic determinism and the limited relevance of transcriptomic or phenomic data collected from leaf tissue at flowering time. Disease related traits have the highest predictive accuracy using expression data, suggesting that gene expression profiles may capture responsive signatures to biotic stress that are detectable by the models and enhance prediction accuracy.

### Transcriptomic features are stronger predictors than genomic markers and capture distinct biological information

To better understand which input features contribute most to phenotype prediction, we asked whether individual gene expression values or genetic markers are more likely to be assigned high importance in our predictive models. We focused on multi-omics models (G + T) that integrate genomic and transcriptomic data and analyzed the top 5% of weighted features for each trait. To evaluate feature contribution, we tested for enrichment of each feature type using fisher’s exact test for gene expression vs. genetic marker among the top-ranked features. For every phenotype, we found that transcriptomic features were significantly more likely to appear in the top 5% of weighted predictors than genomic markers (Figure 4a). The relative proportion of expression vs genetic features varied by trait, but was highest for flowering time and vegetative traits. When looking at feature enrichment in the models that have genetic, transcriptomic, and phenomic data, there were no phenomic features that appeared in the top 5% of weighted features. This is not particularly surprising considering that there are ∼900 phenomic features compared to ∼30,000 genomic and transcriptomic features respectively. However, this disparity does not necessarily imply that phenomic features are uninformative but instead suggests that genomic and transcriptomic datasets contain many more variables of individually smaller effect, while phenomic traits may be fewer in number but more integrative and biologically complex. In this sense, a single phenomic measurement, such as leaf color, may capture the cumulative influence of thousands of underlying genes and transcripts.

**Figure 4.**
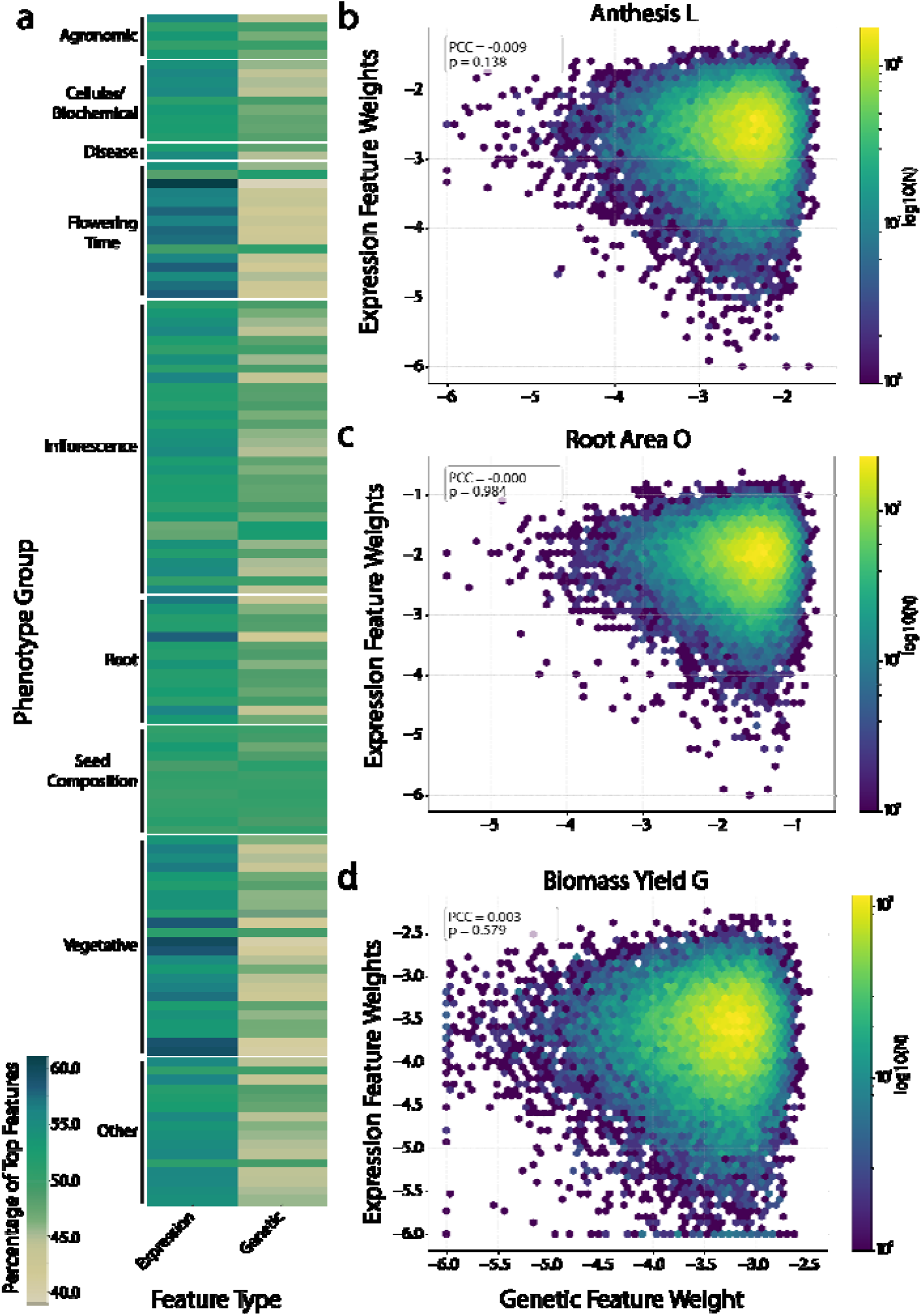
Feature enrichment and overlap in multi-omics predictive models. (a) Heatmap showing enrichment (Fisher’s exact test p-values) of transcriptomic versus genomic features in the top 5% of weighted features across all phenotypes in G + T models. Expression features were consistently more likely to appear among top-weighted predictors. (b–d) Hexbin plots comparing feature importance scores of matched gene regions between genomic and transcriptomic models for three representative phenotypes: (b) anthesis, (c) root mass, and (d) biomass yield. Markers in the top 5% of the genomic model were mapped to the closest gene. The Pearson correlation coefficient (r) and associated p-values are shown in each panel.

To investigate whether the genetic and transcriptomic input spaces contribute redundant or biologically distinct information, we looked at the overlap in the top 5% of features between the expression and marker data. To compare the two inputs, we mapped each of the genomic markers in the top 5% of features to all the nearest genes within a 50kb window to account for any linkage disequilibrium that might cause a marker to be highly weighted even if it is not within the causal gene. We then assessed the overlap between the marker linked gene set and transcriptomic features using a hypergeometric test with a background set of genes, the full set of features. Across all phenotypes, we observed no significant correlation between feature importance scores from the two omics datasets (hypergeometric test, all p-values > 0.05; Figure 4b-d). These results indicate that genomic and transcriptomic datasets capture largely non-overlapping signals, reinforcing the value of integrating multiple data types in predictive modeling.

### Cross-environment prediction of flowering time using transcriptomic data

A unique aspect of our analysis is the ability to predict phenotypes measured with the same genotypes across environments and years. Because we collected paired trait and transcriptomic data from the Wisconsin Diversity Panel grown in both Michigan and Nebraska, we were able to test whether gene expression measured in one environment could accurately predict phenotypes observed in a different field setting. This design also allowed us to assess whether combining transcriptomic data across environments enhances model generalizability and helps capture genotype-by-environment (G × E) interactions. More broadly, it provided an opportunity to evaluate whether expression signatures measured under field conditions represent stable regulatory programs that generalize across locations, or if they instead reflect environment-specific, context-dependent patterns of gene regulation.

To test this, we focused on days to anthesis, a flowering time trait that is widely collected and predicted with relatively high accuracy due to its strong genetic basis, while also exhibiting some environmental sensitivity across diverse field conditions. Interestingly, gene expression collected in Michigan did not always yield the highest predictive performance for phenotypes measured in Michigan. For example, when we used only Michigan transcriptomic data, the models achieved their highest accuracy when predicting days to anthesis measured at the Lincoln, Nebraska field site (Figure 5a). However, when Michigan phenotypic data were included in model training, prediction accuracy for Michigan days to anthesis improved substantially, suggesting that the inclusion of site-specific trait data helps align transcriptomic signals with local environmental responses. This cross-environment result supports our initial hypothesis that transcriptomic data can capture stable regulatory programs that generalize across environments after batch correcting for the field location. Notably, our strongest correlation (PCC = 0.8458) came from the combined G + T model predicting the “Anthesis1_L” phenotype, which includes multiple field sites. This reinforces the idea that integrating transcriptomic data across environments can enhance model generalizability and help capture G×E interactions.

**Figure 5.**
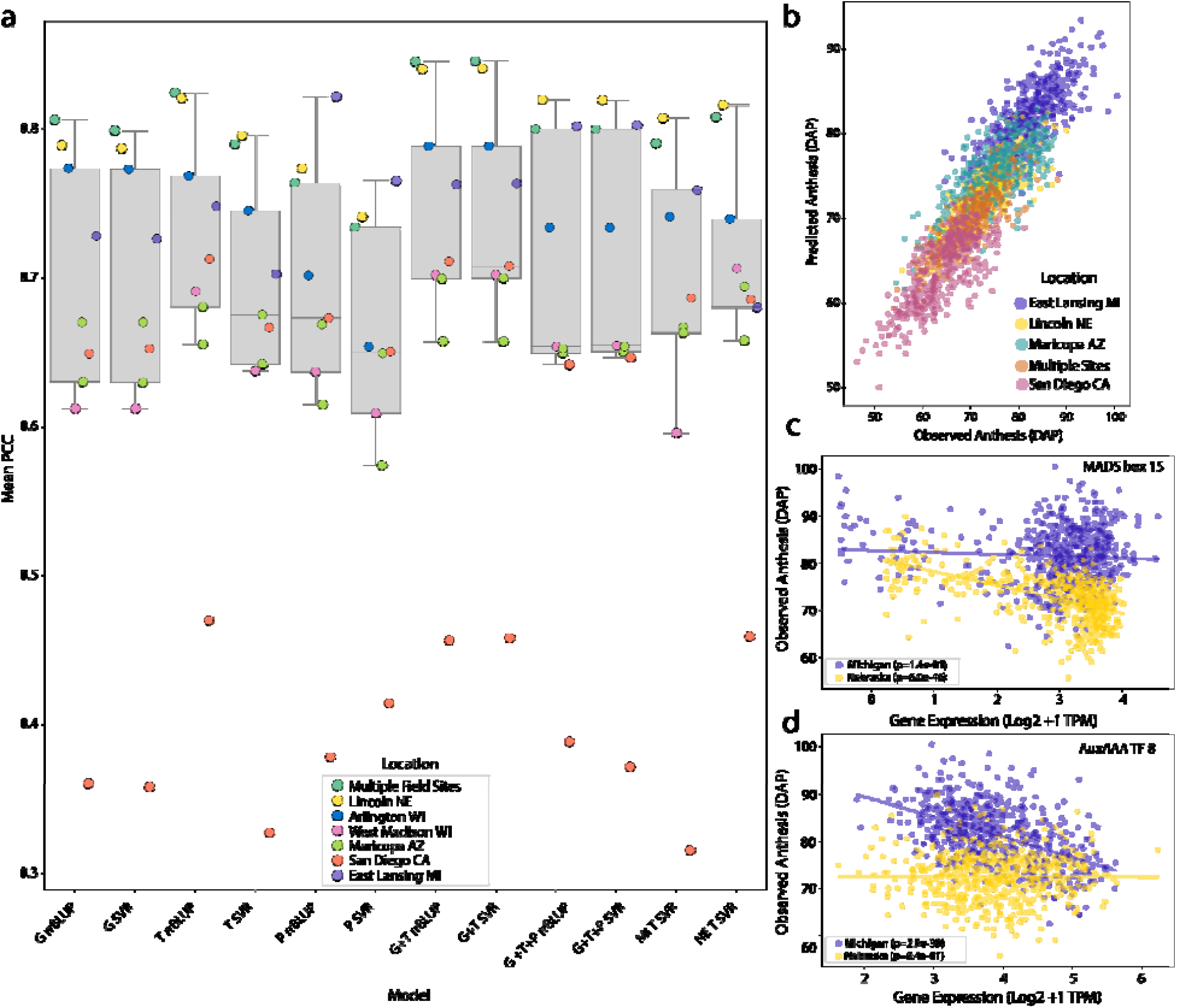
Predicting days to anthesis across environments using transcriptomic data. (a) Predictive accuracy (PCC) of models trained with expression data from Michigan (MI), Nebraska (NE), or both combined (MI+NE) for 16 flowering time phenotypes collected across multiple field trials. Models were trained using SVR with G + T input data. (b) Observed vs. predicted values for anthesis phenotypes across environments using the best-performing SVR G + T model. Points are colored by field site. (c–d) Expression of two highly weighted flowering time-associated genes from the best-performing model plotted against days to anthesis for Michigan and Nebraska phenotypes. (c) MADS-box 15 expression is strongly correlated with anthesis timing in Nebraska but not in Michigan. (d) Aux/IAA transcription factor 8 shows the opposite pattern, correlating with Michigan phenotypes but not those from Nebraska.

This strong performance may reflect the fact that this model was trained on expression data from both Michigan and Nebraska, capturing a generalized or steady-state expression profile that is robust across environments. By incorporating transcriptomic data from plants exposed to different environmental conditions, the model may also be implicitly capturing G × E interactions, allowing it to generalize better to traits measured across diverse field sites. To visualize predictive accuracy across sites, we plotted predicted versus observed values for the anthesis phenotypes, coloring each point by the field location where the phenotype was collected (Figure 5b). These plots reveal a strong correspondence between predicted and observed values, along with a clear ordering of phenotypes across environments. We also compared models trained with expression data from each state individually to those using combined data from both environments. In many cases, using combined data did not consistently improve prediction for traits collected at the same site, suggesting that a core, genotype-driven expression signature may underlie accurate prediction, rather than purely site-specific regulatory responses. This pattern indicates that transcriptomic signal captures underlying developmental programs that are stable across sites, providing predictive power beyond responses tied to the local environment.

To better understand the biological basis of these predictions, we examined the top-weighted features in the best-performing model, SVR G + T (Supplemental Table 1). Two of our top features have been previously associated with flowering time, indicating that our models are capturing biologically meaningful signals present in the expression dataset. However, when we plotted the expression of flowering time genes against days to anthesis in each environment, we observed environment-specific associations. For example, the flowering time associated gene MADS-box 15 ^21^, shows a strong correlation of expression with Nebraska phenotypes but no correlation with expression data from Michigan (Fig 5c). Conversely, expression of another flowering time associated gene, Aux/IAA-transcription factor 8 ^22^, is strongly correlated with anthesis in Michigan but has no correlation with Nebraska anthesis (Fig 5d). These results suggest that while both genes are associated with flowering time, their expression is modulated by the environment, contributing to G × E effects. Although individual gene expression values showed only modest correlations with flowering time, this is expected given the polygenic nature of the trait. High predictive performance arises from the combined effect of many transcripts, not single gene–trait correlations.

## Discussion

Genomic prediction has revolutionized quantitative genetics ^4^, but there is growing interest in leveraging additional datasets representing biological features beyond the genome sequence to improve prediction accuracy ^8^. Our results demonstrate that integrating multi-omic data can enhance predictive power across a broad range of complex traits in maize. Combining genomic markers with gene expression and/or image-derived phenomic features outperformed models based on any single input data type. This aligns with emerging evidence in both plants and animals that incorporating transcriptomic or phenomic information captures variance not explained by DNA markers alone ^12,23^. In our work, the greatest gains came from including gene expression, and models using combined genomic and transcriptomic data yielded higher accuracy for most traits compared to genomic data alone. High-dimensional image features of vegetative indices on their own were generally less predictive than genomes or transcriptomes, but they provided complementary signals for certain traits. For example, adding drone-derived vegetative indices modestly improved prediction of notoriously hard to measure traits like root architecture, suggesting that above-ground imagery can act as a proxy for below-ground phenotypes. These results indicate that incorporating more informative or temporally resolved phenomic data would likely further enhance model performance. These findings reinforce that each data type reflects different biological information including the static genetic potential, the dynamic gene activity in response to conditions, and the integrated organismal phenotype. Harnessing them together can improve trait predictions in ways that any single data type cannot achieve.

Notably, we observed that the benefits of multi-omics integration were trait-specific. No single model or input data type was universally best for all 129 traits, underscoring the importance of trait architecture and utility of different modeling frameworks. Traits with high additive genetic heritability like flowering time had strong predictive accuracy with genomic markers alone, yet even for these traits the inclusion of expression data boosted accuracy further. In contrast, traits strongly influenced by environment or developmental plasticity such as root architecture showed relatively poor predictability with only genomic data, but improved when transcript or image features were included. We also found that a simple linear model rrGBLUP performed on par with a nonlinear machine learning model (SVR) for most traits. This result is consistent with previous comparisons of linear vs. machine learning methods in genomic prediction, which have found no one-size-fits-all algorithm and only modest differences in accuracy in many cases ^7,11,24,25^.

An open question in the field is whether gene expression measurements from one environment can generalize to predict phenotypes in others. Using models built from expression data collected at two locations, we were able to predict traits across nine distinct environments. For example, a model trained with transcriptomic data from plants grown in Michigan predicted days-to-anthesis for plants grown in Nebraska more accurately than it did for Michigan-grown plants. This points to conserved expression patterns within the transcriptomic data, suggesting that our dataset captured core developmental programs that transcend site-specific conditions. By combining expression data from multiple environments, we further improved prediction of flowering time across diverse field sites. This finding is encouraging for breeding applications, as it indicates that a transcriptomic “fingerprint” measured in one context can be leveraged to predict performance in others, potentially capturing genotype-by-environment (G×E) interactions missed by DNA markers alone. Moreover, by examining genes informative for prediction and comparing their expression to phenotypes, we found that different benchmark genes were important in different environments, highlighting a G×E component not detectable using genetic data alone.

By analyzing feature importance, we found that gene expression features dominated multi-omics models, with transcripts consistently overrepresented among the top predictors compared to genetic markers. Unlike discrete genetic variants, transcript abundance represents continuous values that integrate signals from complex regulatory networks and pathways as well as environmental factors, making them strong biomarkers for complex traits. Importantly, genomic and transcriptomic predictors were largely non-redundant, as top SNPs and top transcripts rarely overlapped, showing that each layer contributes unique information. This finding addresses concerns about redundancy in multi-omics prediction and highlights that transcript data provide novel, complementary signals beyond DNA markers. Beyond improving accuracy, these models can also identify key biological pathways underlying trait variation, offering valuable insights for both functional genomics and breeding applications.

Our study demonstrates that integrating genomic, transcriptomic, and phenomic data provides complementary insights and improves prediction of complex traits in maize. This supports the growing shift in breeding and systems biology toward data-rich, multi-layered approaches that capture genetic potential, regulatory networks, and environment-responsive signals. Beyond crop improvement, such models can identify key regulatory pathways, inform synthetic biology and evolutionary studies, and highlight genotype-by-environment interactions not detectable from DNA markers alone. Looking forward, incorporating additional omics layers and temporal data, while balancing cost and scalability, will further advance predictive modeling and accelerate genetic gain in agriculture.

## Methods

### Phenotypic data collection and filtering

Phenotypic data used for model prediction were compiled from two sources (Supplemental Table 2). The first source was a curated dataset from Mural (2022), which collected all previously published GWAS phenotypes using the Wisconsin Diversity Panel ^14^. This initial dataset included 162 quantitative traits sorted into the following categories; vegetative, flowering time, disease, inflorescence, cellular/biochemical, agronomic, root, and seed composition. These phenotypes were collected from 16 different published studies, one of which corresponds to the Nebraska field samples used for RNA-sequencing.

The second trait source consisted of 19 traits measured during the 2021 growing season at the MSU Agronomy Farm, corresponding to the samples collected for RNA-sequencing in the Michigan dataset. A total of 1520 plots were grown in a randomized complete block design with each block consisting of a single plot each of 760 unique genotypes. The traits consist of various agronomic traits including number of leaves, plant height, yield, and other related traits.

Of these 181 original traits, 129 were retained for predictive modeling. Traits were removed if there was low variance (standard deviation < 0.1), high skew (skewness > |1|), and if the trait was conceptually categorical (e.g., tassel openness or branch number), as these characteristics are not well suited to Support Vector Regression modeling.

### Genomic feature processing

A publicly available genetic dataset for the Wisconsin Diversity Panel containing 16,804,001 SNPs and 798 accessions was utilized for the genomic feature space ^15^. Variant filtering and pruning were performed as follows. The raw vcf file was converted to a PLINK ^26^ binary format with the sample ID used for both the family and individual ID. Linkage disequilibrium (LD) pruning was performed using PLINK in sliding windows of 500 SNPs, shifting by 100 SNPs, and removing markers with r2 > 0.2. VCFtools (v0.1.16) ^27^ was used to prune the vcf file to include only variants with a minor allele frequency > 0.1 before being filtered to retain only one SNP per 5 kb. TASSEL (v5.0) ^28^ was used to set a max heterozygosity value of 0.1 to remove any high heterozygous markers in the dataset. This resulted in a final set of 34,153 markers as the genomic input.

### Gene expression feature processing

Gene expression data for the Wisconsin Diversity Panel grown in Nebraska were obtained from ^29^. For the Michigan field trial, gene expression was collected and processed similarly. Five leaf disks were sampled from the pre-antepenultimate leaf (fourth from the topmost fully emerged leaf) within a two-hour window in mid-afternoon. Samples were flash frozen in liquid nitrogen, stored on dry ice, and transferred to a −80°C freezer. Frozen tissue was ground without buffer using a TissueLyzer II (Qiagen; 85300) at 25 Hz in two 30-second intervals, with a 1-minute rest on dry ice between grindings.

RNA was extracted using the MagMax Plant RNA Isolation Kit (ThermoFisher; A47157) on a Kingfisher Flex robot (ThermoFisher; 5400630). Twelve samples per 95-sample batch were run on a 1% agarose gel to verify RNA integrity. Concentrations were quantified using the Quant-IT Broad Range RNA Assay Kit (ThermoFisher; Q10213) on a CLARIOstar Plus plate reader (BMG LabTech). RNA was sent to Psomagen (Rockville, MD) for mRNA purification, cDNA synthesis, and library preparation using Illumina TruSeq strand-specific kits. Libraries were pooled and sequenced on a NovaSeq 6000 (2 × 150 bp), targeting 20 million fragments and 6 Gb per sample.

Thirty-five Michigan genotypes contaminated with ribosomal products were removed from downstream analyses. Raw reads were trimmed using Trimmomatic (v0.33) ^30^, and gene expression was quantified in TPM using Kallisto (v0.46) ^31^ with the primary transcripts from the B73_RefGen_V5 genome ^32^. The initial TPM matrices contained 44,303 genes for Michigan and 39,756 genes for Nebraska. Genes with zero variance, non-expression, or low expression were filtered out, resulting in 33,778 genes for Michigan and 30,302 genes for Nebraska. Raw TPMs were log2-transformed (log2 + 1) and batch-corrected using PyCombat (v0.3.3) ^33^ by field site to account for technical differences between sequencing platforms. This batch-corrected TPM matrix was used as the final transcriptomic feature set for predictive modeling.

### Phenomic Feature processing

Phenomic data were collected by the Remote Sensing and Geographic Information System’s department at MSU in 2021. Twelve flights were conducted between 29 and 126 days after planting. Multispectral imagery was captured from a DJI aerial system at a height of 50m, resulting in a pixel resolution of 4.5cm. The average red, green, blue, red edge, and near infrared reflectance value of each plot were extracted and processed into 10 VIs (Supplemental Table 3). Daily values between flights were interpolated by fitting a locally weighted scatterplot smoothing (lowess) regression for each plot using statsmodels (Seabold and Perktold, 2010).

### Model Training and Testing

Each of the 726 genotypes from the Wisconsin Diversity Panel with multi-omics datasets was assigned to one of five folds such that each fold contained 20% of the total dataset. To protect against overfitting these fold assignments were then used to assign the same genotypes in the same folds for the 751 genotypes in the Nebraska gene expression dataset, filtering each dataset so that each state contained the same genotypes from the Wisconsin Diversity Panel. This ensured that there was never a case where the same genotype had one sample in the training set and one in the testing set. These fold assignments were consistently used in all of the 5-fold cross-validation splits for all the models regardless of input type; genomic, transcriptomic, or phenomic.

For every phenotype in the dataset a separate model was trained and tested, thus there are 129 unique models. According to the assigned labels 5-fold cross-validation was utilized such that each fold was used as the test set once and the remaining 4 folds were used for training.

Then the label was set by iterating through every column in the phenotype dataset and filtering out any missing data from studies that might have excluded genotypes from their analysis or only used a subset of the Wisconsin Diversity Panel. The rrBLUP models were trained and tested in R (v.4.3.3) using the rrBLUP package (v.1.15.0) with the mixed.solve function ^34^. To see whether a machine learning approach would have better predictive accuracy we tested several machine learning algorithms including Random Forest, Support Vector Regression, and XGBoost. We used Support Vector Regression (SVR) moving forward as a comparative nonlinear algorithm because it had the highest performance among those tested. The machine learning models were trained and tested in Python using the SVR model fit with the linear kernel from the package Scikit-learn (v.1.4.1) ^35^.

There were several different separate feature inputs that were utilized. To establish a baseline prediction accuracy the first 75 principal components (PCs) of the genetic data were used to evaluate how well predictions can be made using PCs as a proxy for population structure. The number of PCs used for training and testing was chosen based on the results of a previous machine learning maize paper ^11^. The genomic feature set (G) consisted of the full SNP matrix of 34,153 markers, with one row per genotype. Transcriptomic features (T) consisted of batch corrected expression data from both the Michigan and Nebraska field experiments into a single dataframe separately for a total of 28,221 genes for 577 genotypes. We used 950 phenomic features (P) consisting of vegetative indices extrapolated from drone flyovers during the 2021 field season. In addition, multi-omics models were constructed by combining G and T, with expression values averaged between the genotypes across states, as well as models incorporating G, T, and P together to evaluate predictive performance using all available input spaces.

### Performance Evaluation

For each phenotype, predicted values were plotted against observed values, and fitted with an PCC value to allow comparison across the traits and models. Since 5-fold cross-validation was utilized in the training and testing of the models there are five PCC values for each model and an average was taken to summarize performance. We use PCC as our model evaluation metric, as it is commonly employed in the literature and comparable across diverse trait ranges. These PCC values were then plotted against the heritability values reported by Mural (2022) to assess if heritability is correlated with prediction accuracy. To assess how PCC changes with reported heritability we calculated this ratio for each of the phenotypes. Then each PCC/h2 was plotted as a point for each of the models in a violin plot to assess whether some models predicted traits with higher or lower reported heritabilities (Supplemental Figure 2).

### Feature Weight Investigation

To evaluate which features had the highest impact on the final prediction the marker effects were calculated using the mixed.solve function from the R rrBLUP package. These features were then accumulated into a dataframe to the phenotype they were associated with. This dataframe was then filtered and the highest feature weights were evaluated according to absolute weight to account for features that had a large negative score but greatly impacted the final output.

To assess whether the genetic and expression datasets contained redundant information for prediction we looked at the overlap of highly weighted features. To compare genetic data, represented as SNPs, to expression data we mapped each of the markers to every gene within 50kb using BEDTools ^36^. This was to account for the possibility that a marker is in linkage disequilibrium with a gene of interest and would thus map to a gene that is not associated with the trait of interest. Then the top 5% of features based on absolute value of effect size from each set were assessed for overlap for each phenotype. Then a hypergeometric test using the package scipy.stats was conducted to evaluate whether the overlap between the top weighted features was significantly more than random chance. The feature weights for all of the transcriptomic features and all of the genetic features, assigned to the nearest gene, were then transformed using log10 and plotted against each other in hexbin plots. To assess whether a particular type of feature is enriched in the multi-omics models we performed a Fisher exact test using the package scipy.stats. Then for each phenotype the enrichment of each feature type was plotted in a heatmap.

## Acknowledgements

We thank members of the VanBuren lab for their helpful feedback and discussion. This work was supported in part by the United States Department of Agriculture National Institute of Food and Agriculture (USDA-NIFA) award no. 2022-67013-36118 to R.V. and A.T., and USDA-NIFA, award no. 2020-67014-30902 to A.T., the National Science Foundation (NSF, grant no. OISE-2434687) to R.V. and A.T. and DBI-2213983 to R.V., and by the Advanced Research Projects Agency–Energy (ARPA-E, award no. DE-AR0001367) to J.S. MC was supported by the predoctoral training award T32-GM110523 from the National Institute of General Medical Sciences of the NIH, and BW was supported by the National Science Foundation Research Traineeship Program (NSF-NRT 1828149).

## Data availability

All code used to generate results in the following Github repository: https://github.com/spaddys/Maize_Prediction_Project Note this does not include any of the input datasets as they were too large to upload to Github.

