## Supplemental Figures/Tables for "Predicting complex phenotypes using multi-omics data in maize"

**Supplementary materials**

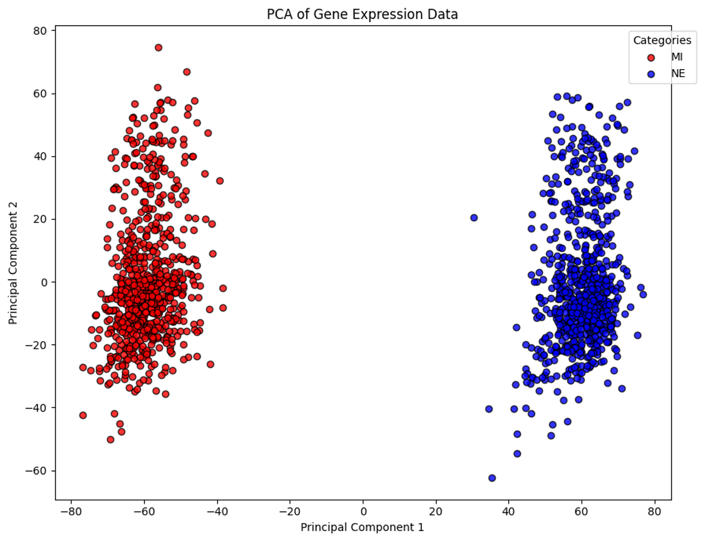

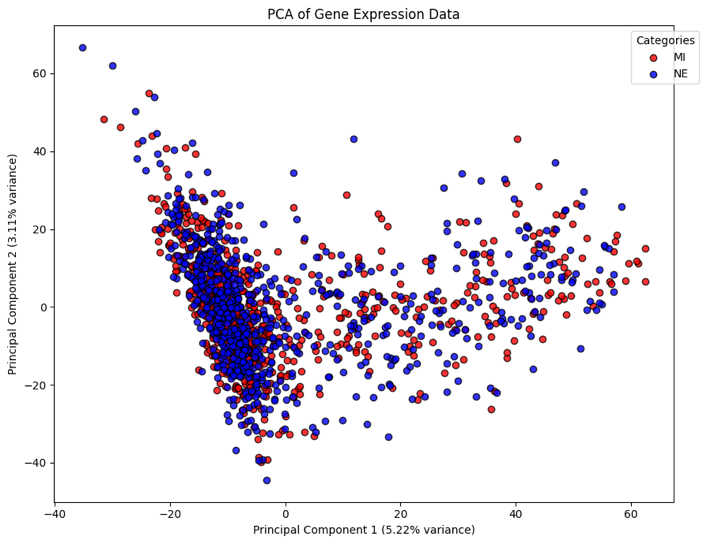

**Supplemental Figure 1. Dimensionality reduction of gene expression input datasets pre and post batch correction.** Principal component analysis (PCA) of the transcriptomic datasets. The left PCA is pre batch correction colored by field location. The right is post batch correction with field location set as the batch variable.

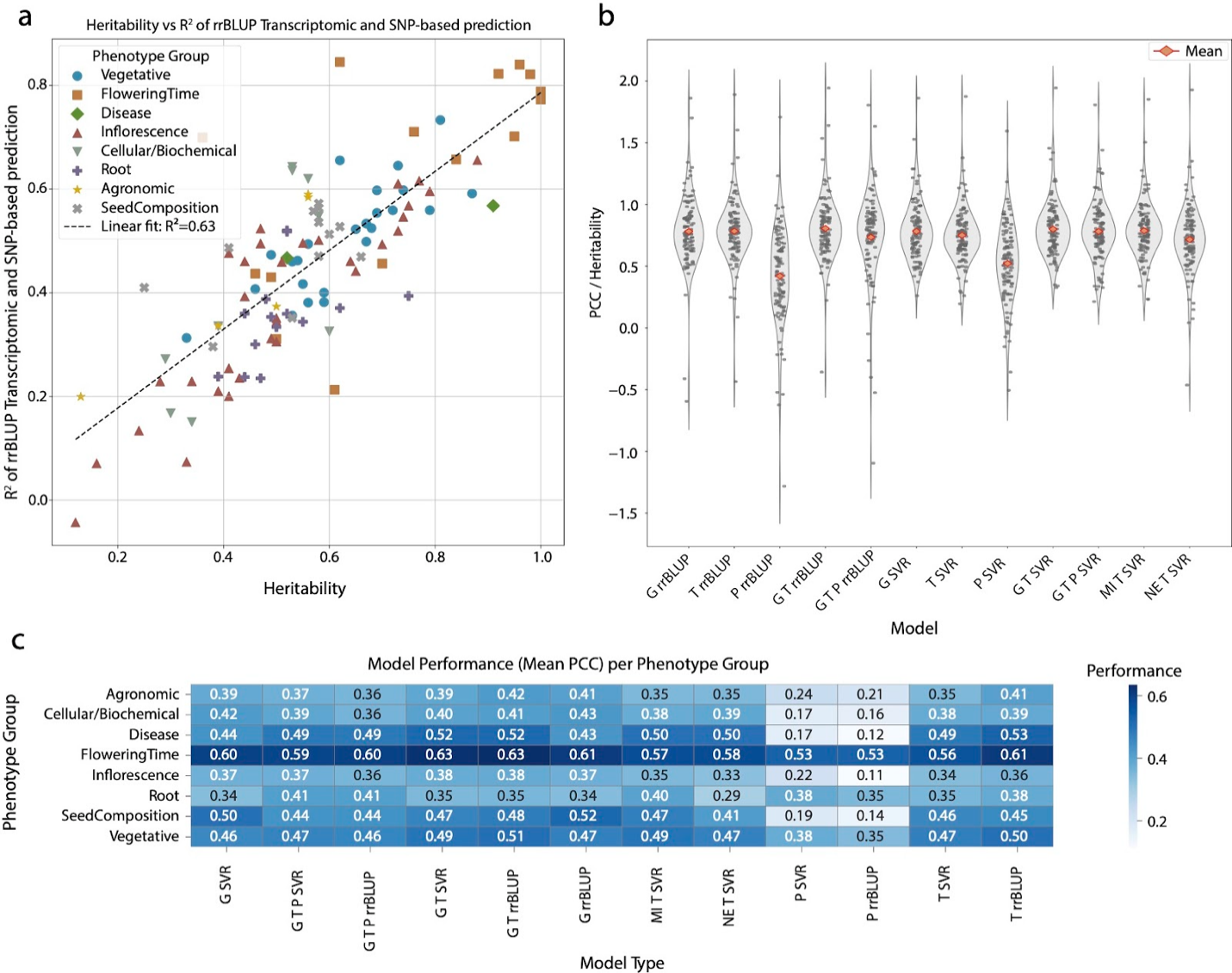

**Supplemental Figure 2. Relationship between PCC and Heritability across input and model types.** Violin plots of the PCC/h2 ratio plotted for every input and model type combination.

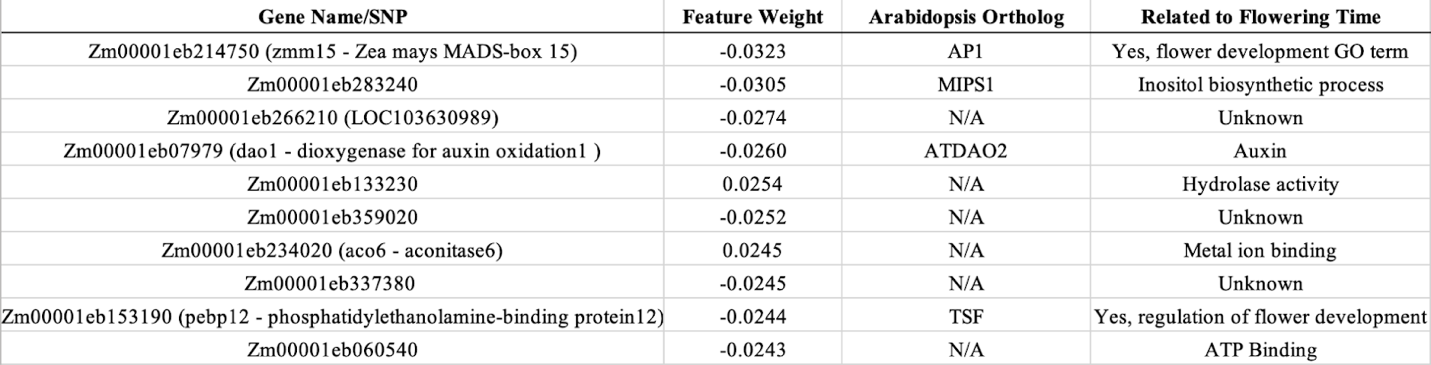

**Supplemental Table 1. Highest weighted features from the best-performing model, SVR G + T for Days to Anthesis.** Highest weighted features according to feature weight absolute value with their respective weight, Arabidopsis ortholog, and any related GO term.

**Supplemental Table 2. Summary of phenotypic data used in this study. (see external excel file).**

| **Index Name** | **Formula** |
| --- | --- |
| **Normalized difference vegetation index - NDVI** | $NDVI = \frac{NIR - Red}{NIR + Red}$ |
| **Green normalized difference vegetation index- GNDVI** | $GNDVI = \frac{NIR - Green}{NIR + Green}$ |
| **Renormalized difference vegetation index - RDVI** | $RDVI = \frac{NIR - Red}{\sqrt{NIR + Red}}$ |
| **Non-linear vegetation index - NLI** | $NLI = \frac{{NIR}^{2} - Red}{{NIR}^{2} + Red}$ |
| **Modified Simple Ratio – MSR** | $MSR =\frac{\frac{NIR}{Red}-1}{\sqrt{\frac{NIR}{Red}} + 1}$ |
| **Chlorophyll vegetation index - CVI** | $CVI = \frac{{NIR}^{2}}{{Green}^{2}}$ |
| **Normalized difference index - NDI** | $NDI = \frac{RedEdge - Red}{RedEdge + Red}$ |
| **RedEdge normalized difference vegetation index - NDVI RedEdge** | $NDVI RedEdge =\frac{NIR - RedEdge}{NIR + RedEdge}$ |
| **Plant senescence reflectance index - PSRI** | $PSRI = \frac{Red - Blue}{RedEdge}$ |
| **RedEdge chlorophyll index – CI RedEdge** | $CIR RedEdge= \frac{NIR}{RedEdge}-1$ |
| **MERIS terrestrial chlorophyll index - MTCI** | $MTCI=\frac{NIR - RedEdge}{RedEdge - Red}$ |

**Supplemental Table 3.** Vegetative Indices used in this study calculated from mean plot level reflectance values of the Red, Green, Blue, Near-infrared (NIR), or Red edge regions.
